# Digging deeper into pain – an ethological behavior assay correlating well-being in mice with human pain experience

**DOI:** 10.1101/2023.08.18.553862

**Authors:** Luke A. Pattison, Alexander Cloake, Sampurna Chakrabarti, Helen Hilton, Rebecca H. Rickman, James P. Higham, Michelle Y. Meng, Luke W. Paine, Maya Dannawi, Lanhui Qiu, Anne Ritoux, David C. Bulmer, Gerard Callejo, Ewan St. John Smith

## Abstract

The pressing need for safer, more efficacious analgesics is felt worldwide. Pre-clinical tests in animal models of painful conditions represent one of the earliest checkpoints novel therapeutics must negotiate before consideration for human use. Traditionally, the pain status of laboratory animals has been inferred from evoked nociceptive assays which measure their responses to noxious stimuli. The disconnect between how pain is tested in laboratory animals and how it is experienced by humans may in part explain the shortcomings of current pain medications and highlights a need for refinement. Here, we survey human chronic pain patients who assert that everyday aspects of life, such as cleaning and leaving the house, are affected by their on-going level of pain. Accordingly, we test the impact of painful conditions on an ethological behavior of mice, digging. Stable digging behavior was observed over time in naïve mice of both sexes. By contrast, deficits in digging were seen following acute knee inflammation. The analgesia conferred by meloxicam and gabapentin was compared in the monosodium iodoacetate knee osteoarthritis model, meloxicam more effectively ameliorating digging deficits, in line with human patients finding meloxicam more effective. Lastly, in a visceral pain model, the decrease in digging behavior correlated with the extent of disease. Ultimately, we make a case for adopting ethological assays, such as digging, in studies of pain in laboratory animals, which we believe to be more representative of the human experience of pain and thus valuable in assessing clinical potential of novel analgesics in animals.

## 1. Introduction

Chronic pain, resulting from inflammation, tissue damage or injury, is a global problem, with patients experiencing reduced quality of life ranging from social isolation, physical incapacity, and mental health disorders [7,28,38]. The issue is exacerbated by the lack of safe and effective medications to provide pain relief. Therefore, development of novel analgesics, particularly those targeted to disease-specific pain mechanisms, has been identified as a priority [33,46]. However, drug discovery efforts for novel analgesics are plagued by a high attrition rate, particularly during transition from pre-clinical testing, using animal (mostly rodent) models, to clinical trials [24]. The disconnect between how pain is assessed in non-verbal rodents and experienced by humans may offer some explanation for this [29,43]. Indeed, several clinical trials of putative painkillers have failed, despite showing promise in preclinical studies [47].

Historically pain in rodents has been inferred from nociception: the neural encoding of noxious stimuli. During such tests, animals are subjected to noxious stimulation, commonly heat or mechanical pressure. The latency to a response (e.g., withdrawal or vocalization), or frequency of nocifensive behaviors (flinching, licking, etc.) are then used to quantify nociception [31]. Animals experiencing pain due to an experimental disease model (e.g., osteoarthritis) are expected to exhibit quicker or more robust reactions. The analgesic potential of new compounds thus relies on animals being able to tolerate noxious stimulation for longer. However, such evoked approaches are not representative of how humans experience, or measure, their pain. Furthermore, these assays are dependent on being able to probe sites affected by pain, limiting their usefulness for certain conditions [27].

Chronic pain patients, living with various conditions, often report an inability to engage with everyday aspects of human life [7,17], which makes a compelling case for studying how ethological animal behaviors are affected by pain models. Indeed, non-reflexive behavioral assays, such as facial grimacing [30], dynamic weight bearing [40], nest building [22], burrowing [45], automated home-cage analytics [20], and machine learning approaches to classify spontaneous behavior [5] are gaining traction as methods to track pain progression and assess analgesic intervention. Not only do these assessments correlate better with how pain affects humans, animals benefit from behaving freely during testing (often in a home-cage environment), and thus such approaches also represent a substantial refinement of the use of animals in pain research.

Another ethological behavior that can be used to assess animal welfare is digging. Distinct from burrowing, which relies on animals being provided with a substrate-filled apparatus, or marbles [10], digging captures the inquisitive nature of mice as they explore an open, uniform environment comprising of compressed digging substrate [12]. Compared to burrowing, digging is quicker to test and less task-driven, which removes a confounding influence on an animal’s motivation to dig. We have already shown digging deficits in acute inflammatory and joint degeneration pain models that correlate with neuronal hyperexcitability [1,8,9], and here sought to further investigate the utility of digging as a readout of animal welfare by focusing on the robustness of naïve animals’ behavior over extended periods of time, as well as the impact of different more translationally relevant pain models.

## 2. Materials and Methods

### 2.1. The Impact of Chronic Pain on Daily Life survey

Ethical approval for human research was granted by The University of Cambridge Psychology Research Ethics Committee (PRE.2023.012). Participants were recruited via social media and with the help of numerous charities based in English-speaking countries with a focus on helping those living with chronic pain conditions (see Acknowledgements) over a period of three months. All participants consented to the use of the anonymous data collected for subsequent analyses and dissemination. Participants were surveyed about their diagnoses of chronic pain, use of analgesics and the impact of their pain on daily activities, some demographic information was also collected on an optional basis. A copy of the survey is included in Appendix B. Where respondents provided optional additional information about medication regimes and non-pharmacological therapies, responses were assigned to categories determined during analysis. Due to international differences in the legality and accessibility of cannabis and cannabidiol (CBD)-containing products as pharmacological treatments for pain, the response of participants that reported sole use of cannabis/CBD to the question about pharmacological interventions was amended to ‘Other’. Use of cannabis/CBD by respondents who also satisfied the criteria to be counted as using prescription, over-the-counter, or both types of drug is still accounted for in the breakdown of medication classes.

### 2.2. Animals

C57BL/6J mice (Envigo) were housed in groups of up to 5 per cage with *ad libitum* access to food and water. Holding rooms were maintained at 21 °C and operated 12-hour light/dark cycles. Mice of both sexes were used for experiments when aged 10-12 weeks (specific compositions of experimental cohorts are defined in corresponding figure legends). All animal work was regulated in accordance with the United Kingdom Animal (Scientific Procedures) Act 1986 Amendment Regulations 2012 and was approved by the University of Cambridge Animal Welfare Ethical Review Body. Behavioral studies commenced after at least a week of habituation in the animal facility. Animals were acclimatized in behavioral testing rooms for at least 30 minutes each day of testing. The room used, time of testing (2 – 4 hours into the start of the light-cycle of animals’ holding room) and experimenters present were kept constant for each individual study to reduce confounds on observed behavior.

### 2.3. Complete Freund’s adjuvant model of inflammatory joint pain

Following induction of anesthesia (ketamine, 100 mg/kg and xylazine, 10 mg/kg, i.p.) and complete loss of consciousness, intra-articular injections were made through the patella tendon using a Hamilton syringe and 30G needle. 10 µl complete Freund’s adjuvant (CFA, 10 mg/ml, Chondrex) or saline was injected unilaterally (ipsilateral knee randomly determined). Knee joint width was measured with digital Vernier calipers before and after injections at specified time points.

### 2.4. Monosodium iodoacetate model of osteoarthritis

Intra-articular injections of 10 µl monosodium iodoacetate (MIA, 0.1 mg/µl, Sigma-Aldrich) or saline were made under inhalation anesthesia (isoflurane: 4% induction, 2% maintenance) unilaterally to a randomly determined knee. Mice were monitored daily for weight loss and knee inflammation. 10 days after injections, following behavioral assessments, mice were injected with gabapentin (30 mg/kg, i.p., Merck) or meloxicam (Metacam, 5 mg/kg, i.p., Boehringer Ingelheim) and behaviors re-tested 90 minutes later.

### 2.5. Dextran sulfate sodium model of colitis

After capturing baseline behaviors and weights, mice were provided with drinking water containing 3% or 1.5% (w/v) dextran sulphate sodium (DSS, Alfa Aesar) for 5 days, after which normal drinking water was provided again for a further 2 days. A control cohort maintained on normal drinking water was run in parallel. Mice were monitored daily: a disease activity index was scored for each animal based on weight loss (0, none; 1, 1-5%; 2, 5-10%; 3, 10-15%, 4, >15%), stool consistency (0, normal; 2, loose stool; 4, watery diarrhea) and presence of blood in stool (0, none; 2, slight bleeding; 4, gross hemoccult). Daily water consumption was also monitored.

### 2.6. Digging behavioral paradigm

Following acclimatization to the behavior room and experimenters, mice were individually transferred to testing cages (49 x 10 x 12 cm) set up with ∼4 cm of tightly packed fine-grain aspen midi 8/20 wood chip bedding (LBS Biotechnology), up to five cages were tested simultaneously. Mice were allowed 3-minutes to explore testing cages without interference, under video surveillance (iPhone 6S camera, Apple). Training sessions were carried out the day before baseline behaviors were assessed, to allow mice to gain familiarity with the set up. Training consisted of mice being placed in test cages as per a standard test. On the rare occasion that an animal did not dig, the individual was placed into a test cage with another animal (from the same home cage) that did meet this criterion, and a second training dig was conducted after a 30-minute break. Separate cages were used to test female and male mice, although when both sexes were studied, both were present in the behavior room for the duration of testing. Following testing sessions, the number of visible burrows remaining at the end of the three-minute test was recorded. Where the same test cage was used multiple times on the same day, half of the digging substrate was refreshed between testing individual mice. Digging behavior was further analyzed offline via video playback at the conclusion of each study, following blinding of all videos. The latency of mice to begin digging was noted and the total time mice spent digging was scored independently by at least two observers; since scores were well correlated (Average R^2^ = 0.7811 ± 0.03 from an average of 34 ± 9 videos scored per experimenter pairing) an average is reported as the time spent digging. We define digging behavior as the vigorous disturbance of digging substrate with all four limbs, which usually begins with a defined planting of the hind legs in a wide stance, followed by rapid movement of substrate under the body using first the fore limbs, and concluding by kicking of the substrate backwards using the hind limbs to produce a burrow.

### 2.7. Automated positional tracking

The open-source Python package Annolid (github.com/healthonrails/annolid/) was used to track the location of mice in videos obtained during assessment of digging behavior. First, a 3-minute video of 5 naïve mice, was used to train a model to identify and track mice in the digging experimental set up. 100 randomly selected frames were manually annotated, model development and Mask R-CNN segmentation was performed using Dectron2 (github.com/facebookresearch/detectron2). The final model had a mean average precision for segmentation of 66.9%. Batch inference was then performed on all videos (5,400 frames each) obtained from DSS colitis studies, to identify the position of each mouse every 10 frames (∼0.3 s), the resulting tracked .csv files were further analyzed in R. Pixel coordinates with a confidence score of >95% were taken forward for analyses, further validation was performed by setting boundaries (300 x 1,150 pixels) for each test cage. Finally, only videos with more than 1 minute worth of tracking were taken forward for analysis, no statistical difference in the duration of validated tracking was observed when comparing on the basis of experimental group (F (2, 3395) = 0.367, *p* = 0.694) or time (F (4, 17566) = 0.95, *p* = 0.441). The standard deviation in x-(α_x)_ and y-position (α_y_) of the center-point of each animal were then used to calculate an explorative area, here defined as the product α_x_.α_y_.

### 2.8. Rotarod

A rotarod apparatus (Ugo Basile 7650) was used to assess the motor coordination of mice. Animals were allowed 1 minute to acclimatize to the apparatus on a slow setting (7 revolutions per minute, rpm), before an accelerating program (7-40 rpm over 5-minutes) was started. Mice were first placed on the rotarod after being trained in the digging assay, to gain some familiarity with the apparatus. Tests were video-recorded, mice were removed from the rotarod if they fell, following two consecutive passive rotations or after 6-minutes of the accelerating program, whichever occurred first. Analysis of videos was performed at the conclusion of studies, following blinding, the latency to passive rotation or fall was scored by a single experimenter.

### 2.9. Pressure application measurement

Mechanical sensitivity of the knee joint was assessed using a pressure application measurement device (Ugo Basile). Animals were scruffed by one experimenter and presented to a second experimenter, blind to the animal’s identity. While lightly holding the hind paw, to maintain a slight bend in the knee of the animal, the second experimenter applied gradual force to each knee joint, squeezing each joint medially with the force transducer. The withdrawal threshold was recorded when an animal withdrew the limb being tested, or after 450 g force (the upper limit threshold) was applied. Each animal was tested twice per time point, with a short break between tests, withdrawal force is reported as an average of the two measurements taken at each time point.

### 2.10. Colon histology

After the final behavior assessments of the DSS study, mice were humanely euthanized by cervical dislocation. A segment of colon was resected ∼1cm from the anus and fixed in 4% (w/v) paraformaldehyde (Sigma-Aldrich) for 6-hours before overnight cryoprotection in 30% (w/v) sucrose (Thermo Fisher Scientific). The following day tissue was embedded in Shandon M-1 Embedding Matrix (Thermo Fisher Scientific) and snap frozen in liquid nitrogen. 20 µm sections of colon were collected on a SuperFrost Plus slides (Thermo Fisher Scientific) using a Leica CM3000 cryostat. Sections were stained with alcian blue (1% w/v in 3% v/v acetic acid; Alfa Aesar; goblet cell stain), hematoxylin (0.2% w/v; Sigma Aldrich; nuclei stain), and eosin (0.5% w/v; Acros Organics; counterstain). Slides were mounted with glycerol and imaged using a NanoZoomer S360 (Hamamatsu). Following blinding, images were scored based on histopathology features: inflammation (0, none; 1, increased immune nuclei in lamina propria; 2, confluence of immune nuclei extending to submucosa; 3, transmural extend of infiltrate), crypt damage (0, crypts intact; 1, loss of basal 1/3 crypt; 2, loss of basal 2/3 crypt; 3, entire crypt loss; 4, epithelial surface erosion; 5, confluent erosion) and ulceration (0, none; 1, 1-2 ulcerative foci; 2, 3-4 ulcerative foci; 3, extensive ulceration). At least two sections, separated by 150 µm were scored per colon, sections were scored independently by 3 experimenters, an average of their scores is reported.

### 2.11. Data analysis and statistics

The number of animals and sex composition of studies are detailed in each corresponding figure legend. Recovery Index is defined as (post-drug behavior – pre-drug behavior)/(Baseline behavior – pre-drug behavior), i.e., Recovery Index < 0 represents a worsening of behavior, Recovery Index = 0 signifies no change, Recovery Index = 1 indicates complete recovery, Recovery Index > 1 represents more/ more sensitive behavior. Data are presented as mean ± standard error of the mean. Appropriate analyses were selected according to the number of factors being compared and whether data met the assumptions for parametric analyses, the statistical tests employed are stated in corresponding figure legends. All analyses were performed in R, and *p* values less than 0.05 were considered significant.

## 3. Results

### 3.1. Chronic pain prevents full engagement with daily life

To assess the relevance of using ethological behaviors as a means of assaying nociception in laboratory animals, it was important to first confirm that natural human behaviors are affected in those experiencing chronic pain. To address this, we ran an online survey (Appendix B). Over a three-month recruitment period, 341 participants completed the survey. An optional demographic section (completed by 339 respondents) revealed that the majority (83.2%) of responders identified as female (Fig. S1A), the ages of respondents were relatively uniformly distributed (Fig. S1B), the overwhelming majority self-identified as White (85.8%; Fig. S1C) and most responses came from those living in United Kingdom (53.3%), New Zealand (25.3%) and Australia (11.3%; Fig. S1D). Respondents all confirmed they experience chronic pain (regular bouts of pain persisting at least 3 months) due to a medically diagnosed health condition. The underlying health conditions varied, with the largest numbers experiencing chronic pain due to: fibromyalgia, osteoarthritis, migraine and irritable bowel syndrome (IBS, Fig. 1A; full diagnosis breakdown available in Table S1); 55.1% of respondents had diagnoses of more than one condition causing them chronic pain (Fig. 1B).

Respondents were asked about the frequency and intensity of their pain, with constant, low intensity pain being most commonly reported (27.0%), followed by low intensity pain on a daily basis (19.9%; Fig. 1C). In addition to their usual experience of pain, 97.9% of patients expressed that they also experience flare ups of more intense pain (Fig. 1D). The frequency of flare ups varied from daily to less frequent than monthly (Fig. 1E). Participants were also asked about their use of drugs to manage pain, with 65.7% of respondents using prescription medication, a further 21.4% relying on over-the-counter analgesics, 4.1% using a combination of both prescription medication and over-the-counter drugs and 1.5% of respondents using cannabis/cannabidiol-containing products only (classified as ‘Other’). 7.3% of respondents reported not using any pharmacological intervention for their pain (Fig. 1F). Of the 258 participants that shared details on their use of drugs, the majority (68.6%) use multiple drugs to try and manage their pain (Fig. S1E). A breakdown of medication classes revealed that weak opioids (used by 31.0% of participants that shared details on their drug use) and paracetamol (28.3%) were the most commonly used (Fig. S1F), with one third (33.3%) taking some form of antidepressant (Fig. S1F). Of the 308 respondents using drugs to treat pain, only 5.5% found them to be completely effective at relieving pain, with most (82.4%) finding them somewhat effective and 12.1% not achieving any pain relief with their medications (Fig. 1G). Additionally, 57.5% of respondents reported experiencing unpleasant side effects from taking medication (Fig 1H). Emphasizing the desire of chronic pain patients for effective therapies, 74.2% of patients also make use of non-pharmacological treatments to treat their pain (Fig. 1I), with 62.8% of those using multiple alternative therapies (Fig. S1G). Analysis of the types of non-pharmacological therapies used by participants placed physiotherapy, massage, thermal, and exercise as the most popular non-pharmacological treatments used (Fig. S1H).

Questions probing the effect of pain on daily life, confirmed that pain has a mostly negative impact on many varied aspects of daily life, often taken for granted by those not living in chronic pain. For instance, 63.9% of respondents agreed that tending to their personal hygiene could cause them or increased their pain (Fig. 1J). An overwhelming majority of participants agreed that pain has prevented them from undertaking housework, engaging with physical activity, socializing and leaving the house. Whether pain interfered with non-physical recreational activities (e.g., reading) divided respondents the most, with 56.6% agreeing that pain can prevent them and 29.9% disagreeing. Regarding sleep, 87.7% of respondents reported that their sleep quality is impaired by their pain and 91.8% participants agreed that their pain contributed to low mood. Perhaps most importantly was the finding that 94.1% of participants agreed that they are unable to live life as they would if they did not experience pain. Overall, these data support the notion that pain reduces an individual’s ability and motivation to engage in everyday activities and that it would make sense to examine how equivalent behaviors in animals are affected by models of pain and pharmacological intervention.

**Figure 1.**
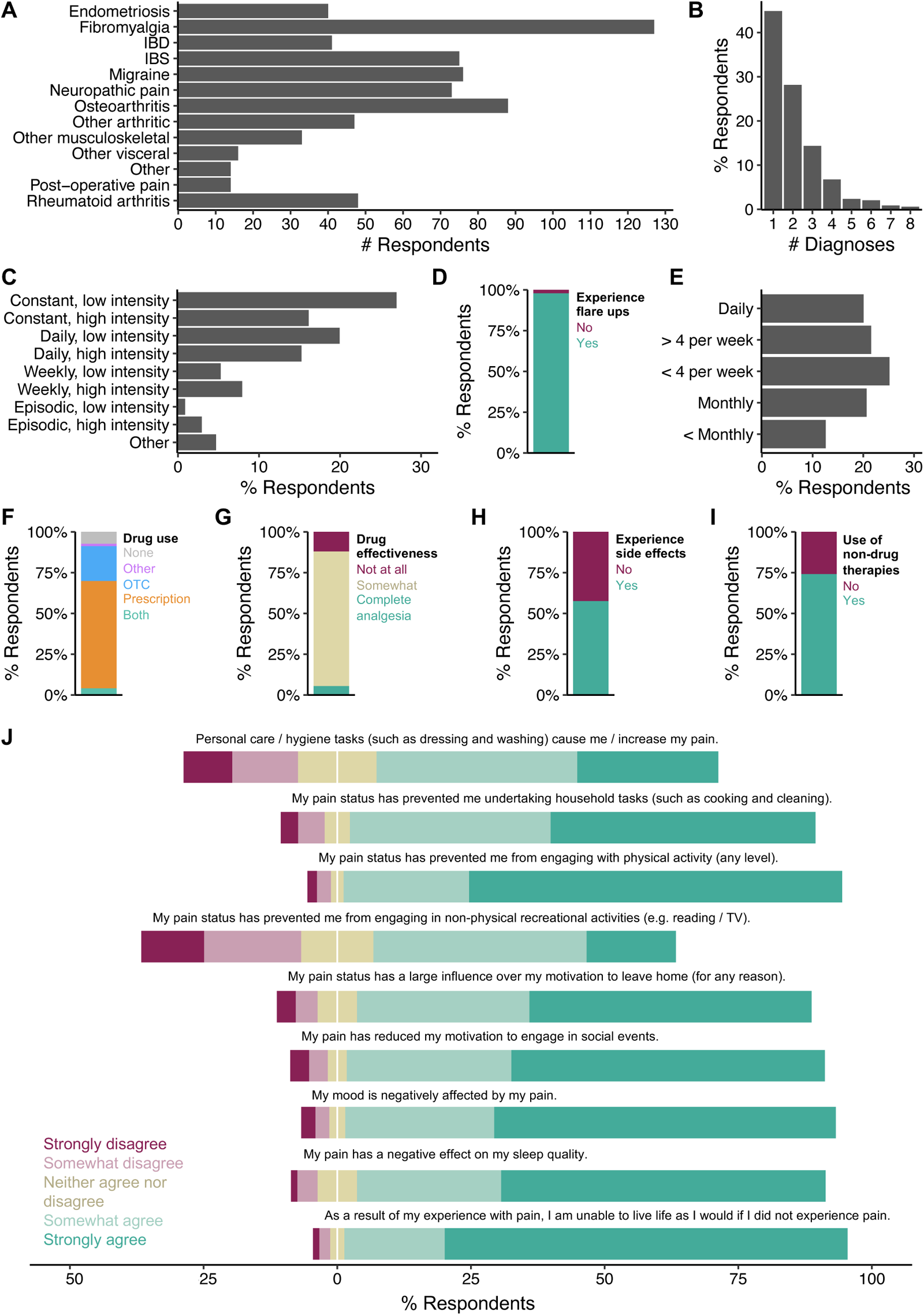
Experiences with pain have a negative impact on the daily lives of human patients living with chronic pain. Summary of results from The Impact of Chronic Pain on Daily Life survey, completed by 341 participants. **(A)** Number of diagnoses of conditions that caused respondents chronic pain. **(B)** Frequency distribution of the number of diagnoses of chronic pain conditions by respondent. **(C)** Frequency and intensity of the pain experienced by respondents on usual basis. **(D)** Proportion of respondents experiencing flare ups of pain, in addition to their usual experience with pain. **(E)** Distribution of the frequency of additional flare ups. **(F)** Use of medications to manage pain, OTC = over-the-counter. **(G)** Effectiveness of drugs to manage pain. **(H)** Occurrence of side effects from pain medications. **(I)** Use of non-pharmacological therapies to manage pain. **(J)** Opinion scales on how experience with pain affects various aspects of daily life.

### 3.2. Digging is a robust measurable ethological behavior

To better understand the stability of digging behavior, naïve mice of both sexes were studied over a period of 10 days, a common timeline for many rodent pain models (Fig. 2A). The latency to dig, time mice spent digging and number of burrows produced during each test period were measured. Additionally, on each test day a random subset of mice was re-tested after a 30-minute break to assess the consistency of digging behaviors. The time taken for mice to begin digging after starting testing remained relatively consistent across the study (F (5,45) = 1.649, *p* = 0.167; Fig. 2B), with an overall average latency to dig of 63 ± 5.5 s. Latencies remained similar across test digs when repeated after a 30-minute break (Test 1, 60.7 ± 17.4 s vs. Test 2, 52.4 ± 15.7 s, t = 0.678, df = 9, *p* = 0.515; Fig. 2C). Similarly, the time mice spent digging did not differ with each time point tested (F (5, 45) = 2.333, *p* = 0.058; Fig. 2D), with mice spending an average of 11.9 ± 1.0 s digging each day. A slight increase in the time mice spent digging was observed during the second time tested, although the difference was not significant (Test 1, 9.67 ± 2.5 s vs. Test 2, 14.5 ± 3.3 s, t = -2.164, df = 9, *p* = 0.058; Fig. 2E). The cohort of naïve mice produced an average of 3.98 ± 0.1 burrows, regardless of when tested during the study (χ^2^(5) = 5.86, *p* = 0.320; Fig. 2F). The number of burrows produced between repeat tests on the same day remained consistent as well (Test 1, 4.4 ± 0.2 vs. Test 2, 3.9 ± 0.2, W = -25, *p* = 0.180; Fig. 2G). In the small cohort tested, the average time male mice spent digging was slightly higher than that recorded for females, although no significant difference was seen (Time digging: F (1, 56) = 3.866, *p* = 0.054; Fig. 2D), importantly, digging durations remained consistent over time when scrutinized by sex. In summary, the digging behavior of naïve mice is stable over timeframes commonly used in behavioral studies assessing pain.

**Figure 2.**
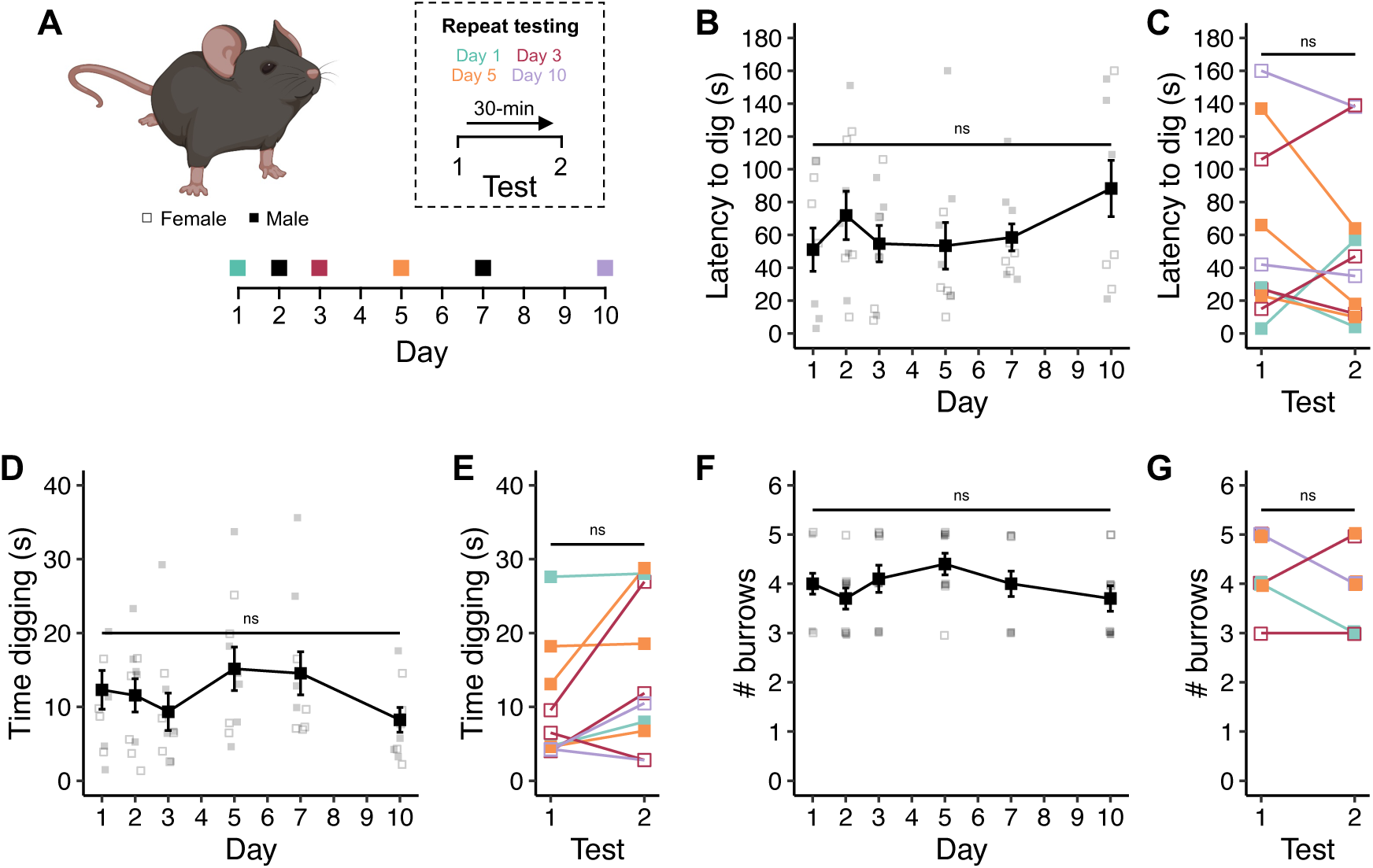
Digging is a robust and measurable ethological behavior. **(A)** Timeline of behavioral experiments. **(B)** Average latency to dig across experimental timeline. **(C)** Latencies to dig of individual mice when tested twice in one day, 30-minute break between tests. **(D)** Average time spent digging across experimental timeline. **(E)** Digging durations of individual mice when tested twice in one day, 30-minute break between tests. **(F)** Average number of burrows dug by mice at each test point. **(G)** Number of burrows dug by individual animals when tested twice in one day, 30-minute break between tests. **(B, D)** Repeated measures ANOVA. **(C, E)** Paired t-test. **(F)** Friedman repeated measures ANOVA. **(G)** Wilcoxon signed-rank test. n = 8, 4 females (denoted by open symbols), 4 males (denoted by closed symbols).

### 3.3 Acute inflammation of the knee joint decrease digging behavior

Injection of complete Freund’s adjuvant (CFA) into the knee joint of mice is a commonly used model of acute inflammatory pain [19]. To confirm that digging behaviors can be used as a readout of inflammatory pain, mice were injected unilaterally through the patellar tendon with either CFA, or saline serving as a control for intra-articular injection (Fig. 3A). Monitoring of joint inflammation revealed a distinct difference between the two cohorts with respect to injection and time (Interaction Time:Injection: F (1, 12) = 30.036, *p* < 0.001), but no apparent sex difference was observed for the development of this model (Sex: F (1, 12) = 0.949, *p* = 0.349). Mice that received CFA exhibited joint inflammation, evident from an increased ratio in the diameter of the ipsilateral joint relative to the contralateral joint (Baseline, 0.99 ± 0.01 vs. 24-hours, 1.29 ± 0.03, t = -9.240, df = 9, *p.adj* < 0.0001; Fig. 3B). No swelling was observed for mice injected with saline (Baseline, 1.00 ± 0.00 vs. 24-hours, 1.01 ± 0.02, t = -0.778, df = 5, *p.adj* = 0.472; Fig. 3B). Consistent with the absence of any swelling, the latency to dig was not affected by intra-articular injection of saline (Baseline, 17.5 ± 7.8 s vs. 24-hours, 12.7 ± 3.9 s, t = 0.520, df = 5, *p.adj* = 0.625; Fig. 3C), whereas mice injected with CFA took longer to start digging 24-hours post-injection (Baseline, 33.1 ± 8.2 s vs. 24-hours, 68.2 ± 14.5 s, t = – 2.857, df = 9, *p.adj* = 0.019; Fig 3C). The time mice spent digging and the number of burrows dug during the test period were also unaffected by saline injection (Duration: Baseline, 34.8 ± 8.3 s vs. 24-hours, 32.7 ± 5.8 s, t = 0.348, df = 5, *p.adj* = 0.742; Fig. 3D; Burrows: Baseline, 4.1 ± 0.5 vs. 24-hours, 4.0 ± 0.4, t = 0.542, *p.adj* = 0.611; Fig. 3E). However, injection of CFA reduced both the amount of time mice spent digging (Baseline, 30.6 ± 5.2 s vs. 24-hours, 7.7 ± 2.0 s, t = 5.061, df = 9, *p.adj* = 0.0007; Fig. 3D) and the number of burrowing sites recorded (Baseline, 4.7 ± 0.4 vs. 24-hours, 2.3 ± 0.5, t = 5.041, *p.adj* = 0.0007; Fig. 3E). Importantly, no adverse effect was seen when mice were tested on the rotarod following CFA-induced inflammation (Baseline, 296.6 ± 34.8 s vs. 24-hours, 354.0 ± 6 s, t = -1.614, df = 4, *p.adj* = 0.182; Fig. S2), indicating that mice retain normal locomotor function. These results thus suggest that the inflammation resulting from intra-articular injection of CFA reduces the motivation of mice to dig, while they likely remain able to. Possible explanations for the observed digging deficit may arise from mice finding digging painful following induction of inflammation, or, the occurrence of spontaneous pain, resulting from the inflammatory insult, triggering an apathetic withdrawal.

**Figure 3.**
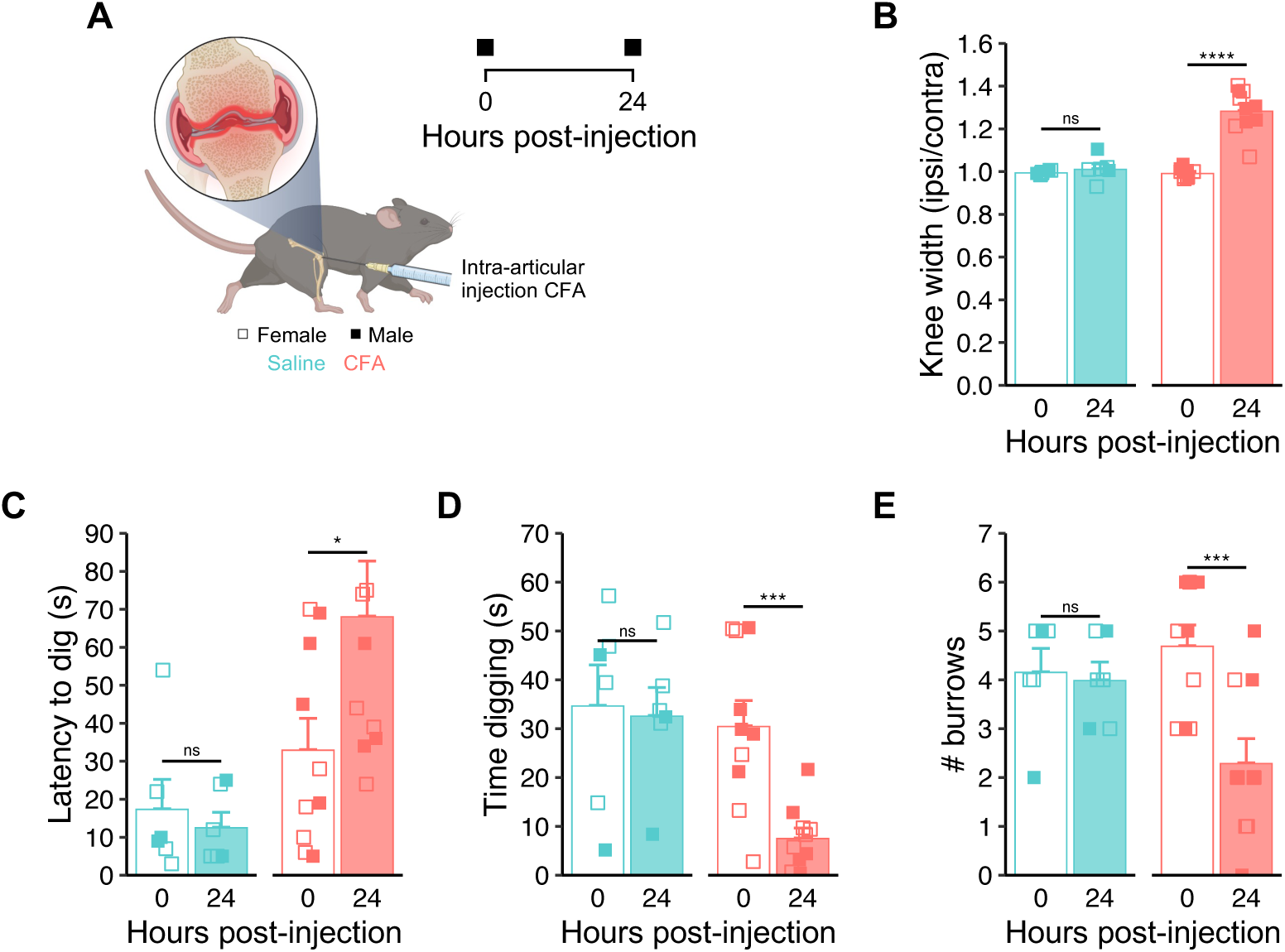
Acute inflammation of the knee joint decreases digging behavior. **(A)** Schematic representation of the complete Freund’s adjuvant (CFA) knee model (created with BioRender.com) and experimental timeline. **(B)** Ratio of ipsilateral knee width to contralateral knee width on the day of injections and 24-hours after. **(C)** Latency of mice to dig**. (D)** Time spent digging before and 24-hours after CFA injection. **(E)** Number of visible burrows at the conclusion of 3-minutes digging test. ∗ *p.adj* < 0.05, ∗∗∗ *p.adj* < 0.001, ∗∗∗∗ *p.adj* < 0.0001: Repeated-measures ANOVA followed by Bonferroni corrected post-hoc. n = Saline: 6, 4 females (denoted by open symbols), 2 males (denoted by closed symbols); CFA: 10, 5 females (denoted by open symbols), 5 males (denoted by closed symbols).

### 3.4 Digging deficits are also seen in a more translationally relevant model of osteoarthritis

Having confirmed that the digging activity of mice is reduced following acute inflammation, the suitability of digging as a welfare readout in a more chronic model was assessed. Injection of monosodium iodoacetate (MIA) into the knee joint of mice results in a transient inflammatory response and progressive joint degeneration, recapitulating many of the aspects of osteoarthritis (OA) [34]. Following assessment of baseline behaviors, mice received a unilateral intra-articular injection of either MIA or saline, and were studied for 10 days (Fig. 4A). Swelling of the injected knee, relative to the contralateral joint, was used to monitor model development with the injection mice received and experimental time having a significant effect (Interaction: Time:Injection: F (3.58, 71.55) = 15.71, *p* < 0.0001; Fig. 4B). The sex of mice was not found to influence development of OA-like symptoms, as inferred from knee swelling (F (1, 20) = 2.33, *p* = 0.142; Fig. 4B). The extent of joint swelling following MIA injection, peaked 24-hours post-injection, but then steadily declined throughout the study (F (10, 220) = 54.2, *p.adj* = 0.035; Fig. 4B). No such swelling was seen following injection of saline (F (10, 220) = 0.127, *p.adj* = 0.999; Fig. 4B). In line with the progressive degeneration of the injected joint following injection of MIA, the latency of mice to dig was observed to increase over time (F (4, 100) = 16.54, *p.adj* < 0.0001; Fig. 4C), while minimal change was seen for saline-injected mice (F (4, 100) = 0.151, *p.adj* = 0.962; Fig. 4C). The time spent digging declined with time following injection of MIA (F (4, 100) = 8.35, *p.adj* < 0.0001; Fig. 4D), as did the number of burrows dug during the test period (F (4, 100) = 16.086, *p* < 0.0001; Fig. 4E), but neither were affected for mice injected with saline (Duration: F (4, 100) = 0.207, *p.adj* = 0.934; Fig. 4D; Burrows: F(4, 100) = 0.231, *p.adj* = 0.92; Fig. 4E). The deficits in digging behavior observed here are likely the result of mice experiencing pain, given that development of mechanical hypersensitivity was observed for the MIA-injected knee joint (F (4, 70) = 13.707, *p.adj* < 0.0001; Fig. 4F). As expected, mechanical sensitivity of knee joints injected with saline and the contralateral joints of both MIA- and saline-injected mice remained constant throughout the duration of the study (Ipsi_saline_: F (4, 70) = 1.1, *p.adj* = 0.363; Fig. 4F; Contra_MIA_: F (4, 70) = 1.89, *p.adj* = 0.122; Fig. S3A; Contra_saline_: F (4, 70) = 0.902, *p.adj* = 0.468; Fig. S3A). The ability of mice to use their joints when required was not affected by the MIA model, as is evident from the lack of an effect on the time either group of mice could stay on the rotarod (MIA: F (4, 100) = 1.833, *p.adj* = 0.128; Fig. S3B; Saline: F (4, 100) = 0.874, *p.adj* = 0.482; Fig. S3B). Considered together, these results are indicative of mice experiencing pain and consequently reduced motivation to explore and to interact with the environment with the onset of arthritis-like symptoms.

A variety of medications including, gabapentinoids, such as gabapentin, and non-steroidal anti-inflammatory drugs (NSAIDs), such as meloxicam, are prescribed to human OA patients, to alleviate the pain associated with their condition[2,48]. In addition, administration of both gabapentin and meloxicam to rodents following progressive degeneration of the knee joint can remedy mechanical hypersensitivity [5,37,44]. Although an improvement in certain aspects of spontaneous behavior can be seen, comprehensive 3D pose analysis has shown that gabapentin does not return the spontaneous behavior of mice to their pre-injury state [5]. Such reports question the analgesic properties of gabapentin, when focusing on metrics more relevant to human pain experience. The effects of gabapentin and meloxicam on the digging deficits induced by the MIA model of OA were thus tested. Following behavior testing on day 10, mice were injected with either gabapentin or meloxicam and behaviors re-assessed 1.5-hours later (Fig. 4A). The level of analgesia conferred by each agent was inferred from recovery indices calculated based on the mechanical sensitivity of the ipsilateral knee joint and digging duration. Whereas both drugs seemed equally effective at relieving evoked pain measured through application of mechanical pressure to the knee joint (Median mechanical hypersensitivity recovery index: GBP, 0.52; MEL, 0.48; Fig. 4G), better remedy of digging duration deficits was seen following meloxicam administration (Median digging duration recovery index: 0.72; Fig. 4G) compared to gabapentin (Median digging duration recovery index: 0.00; Fig. 4G). Gabapentin is known to be a sedative [14], which may help explain these findings, indeed, mice struggled more on the rotarod test 1.5-hours after receiving gabapentin (Median % change in time on rotarod: -80.9%; Fig. 4H) than meloxicam (Median % change in time on rotarod: -5.0%; Fig. 4H). Importantly, these findings are consistent with the opinions of participants in our survey, living with OA, who reported use of gabapentinoids (Fig. 4I) and NSAIDs (Fig. 4K) to manage their pain. An overwhelming majority of those using gabapentinoids (75%) felt that their medication regime was only partially effective at reducing their pain, with the remaining 25% reporting a complete lack of effectiveness (Fig. 4Ji). On the other hand, 8% of OA patients regularly taking NSAIDs reported achieving complete pain relief, with 88% finding some benefit from taking NSAIDs and just 4% respondents reporting no pain relief (Fig. 4Li). Additionally, more gabapentinoid users (68.75%) reported experiencing unpleasant side-effects from their use of pain medications (Fig. 4Jii), compared to 48% of OA patients relying on NSAIDs for pain relief (Fig. 4Lii). The questionable analgesic properties of gabapentin in the digging assay reported here, corroborates the lack of effectiveness reported by human patients who use similar medications, evident from our own survey (Figs 4I-L) and the findings of others [13]. Considering both gabapentin and meloxicam reversed evoked pain assay deficits, our data only strengthen the notion for the use of ethological approaches in preclinical tests of novel analgesics.

**Figure 4.**
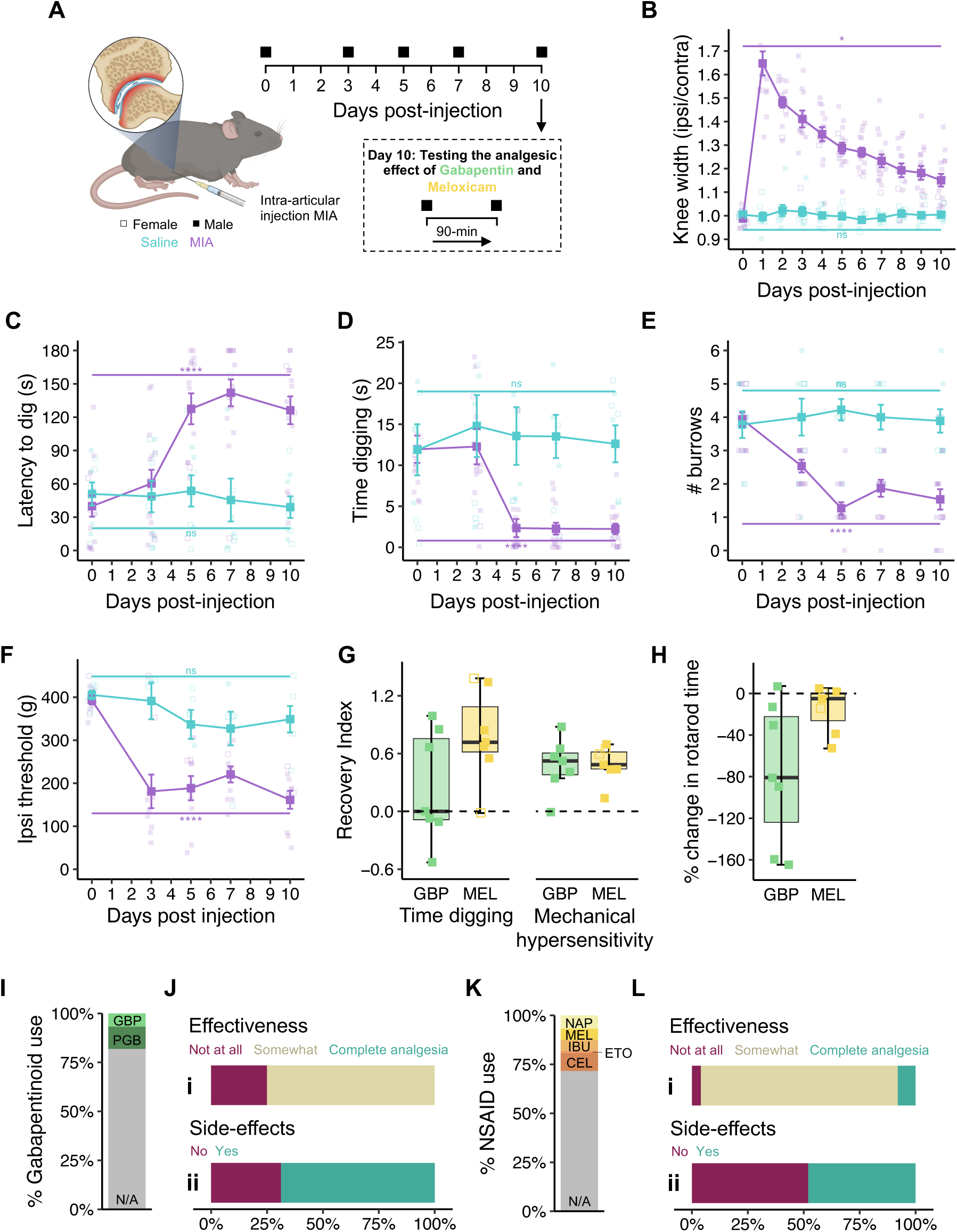
Digging behavior is decreased in the MIA model of osteoarthritis. **(A)** Schematic representation of the MIA model (created with BioRender.com) and experimental timeline. **(B)** Ratio of ipsilateral knee width to contralateral knee width. The **(C)** latency to dig, **(D)** time digging, **(E)** number of burrows dug and **(F)** mechanical sensitivity of the ipsilateral knee joint were measured across experimental time. **(G)** Comparison of the recovery in time spent digging and mechanical sensitivity of the knee join following administration of gabapentin (GBP) or meloxicam (MEL). **(H)** Percentage change in rotarod performance following administration of analgesics. **(I)** Proportion of respondents from Impact of Chronic Pain on Daily Life survey using gabapentinoids (GBP = gabapentin; PGB = pregabalin) to manage pain. **(J)** Opinion of survey respondents that use gabapentinoids on the **(i)** effectiveness of and **(ii)** experience of side-effects from their pain medications. **(K)** Proportion of respondents from Impact of Chronic Pain on Daily Life survey using non-steroidal anti-inflammatory drugs (NSAIDs, NAP = naproxen; MEL = meloxicam; IBU = ibuprofen; ETO = etoricoxib; CEL = celecoxib) to manage pain. **(L)** Opinion of survey respondents that use gabapentinoids on the **(i)** effectiveness of and **(ii)** experience of side-effects from their pain medications. ∗ *p.adj* < 0.05, ∗∗∗∗ *p.adj* < 0.0001: Repeated measures ANOVA followed by Bonferroni corrected post-hoc, annotated statistical differences are the effect of time for each experimental group. **(B-F)** n = 2 MIA females (denoted by purple open symbols), 8 MIA males (denoted by purple closed symbols, 5 saline females (denoted by blue open symbols), 4 saline males (denoted by blue closed symbols). **(G-H)** n = 7 males received gabapentin (denoted by green closed symbols), n = 2 females received meloxicam (denoted by yellow open symbols), n = 5 males received meloxicam (denoted by yellow closed symbols).

### 3.5 Digging behavior is predictive of the extent of intestinal inflammation in a visceral pain model

Painful conditions affecting visceral organs are highly prevalent but are difficult to study in both human patients and animal models due to their internal nature. A behavioral readout that offers insight to animal wellbeing in relation to the extent of disease, would thus likely be highly useful. To this end, the effect of an experimental model of ulcerative colitis, a form of inflammatory bowel disease, on digging behaviors was tested. Addition of dextran sulfate sodium (DSS) to the drinking water of rodents triggers colitis-like intestinal inflammation through disruption of the intestinal epithelia [35]. An acute DSS-colitis model, where mice are provided with DSS-containing drinking water for 5 days, before returning to normal water for a further 2 days was used to assess the suitability of digging behavior to infer animal wellbeing in visceral pain (Fig. 5A). The extent of colitis symptoms correlates with the concentration of DSS provided [15], we thus elected to study two doses (3% w/v and 1.5% w/v) with the aim of interrogating whether digging parameters can distinguish between disease severity. A control cohort kept on water for the duration of the study was run in parallel. Upon starting the model, mice were monitored daily to estimate the extent of disease progression, producing a disease activity index (DAI). As expected, a significant difference in the extent of disease activity was seen dependent on the dose of DSS that mice were administered (F (2, 228) = 181.144, *p* < 0.0001; Fig. 5B). In line with the work of other groups using the same model [18], a sex-dependent effect was also observed (F (1, 228) = 32.186, *p* < 0.0001; Fig. 5B), which was most noticeable for the 1.5% w/v dose of DSS (DAI at Day 7: Females, 2.4 ± 0.3 vs. Males, 8.0 ± 0.3, t = -14.00, df = 4, *p.adj* = 0.0002). The DAI data suggest that males are more susceptible to DSS colitis, however, this is not due to higher consumption of DSS-containing water, as the average daily water consumption of animals over the first five days of the experiment was consistent across groups and sexes, when corrected to animal body weight (Interaction: Sex:Group F(2, 24) = 0.03, *p* = 0.971; Fig. S4A).

In line with the progressive nature of the model, a trend for an increased latency to dig was seen for mice that received DSS-containing water, with the latency of mice on the 3% w/v dose increasing 64% from day 0 to day 7. Although the effect of dose on latency was not found to be significant (F (2, 24) = 1.039, *p =* 0.369; Fig. 5C), minimal change (-22%) was seen in the latency of mice kept on regular drinking when comparing day 7 to day 0, suggesting that mice on the highest dose of DSS began to lose in motivation to dig by the end of the study. In agreement with this, a decrease in the time mice spent digging was observed over experimental time (F (2.69, 64.61) = 7.903, *p* = 0.0002; Fig. 5D), which was also affected by sex (F (1, 24) = 4.791, *p* = 0.039; Fig. 5D). These effects were most noticeable at the conclusion of the study, where mice that received the high dose of DSS dug substantially less than the control cohort for both sexes (Female: 0% DSS, 7.6 ± 0.7 s vs. 3% DSS, 2.3 ± 0.6 s, t = 5.43, df = 4, *p.adj* = 0.017; Male: 0% DSS, 6.6 ± 0.5 s vs. 3% DSS, 2.0 ± 0.8 s, t = 6.14, df = 4, *p.adj* = 0.011). However, only males were seemingly affected when comparing mice that received 1.5% w/v DSS to the control group (Female: 0% DSS, 7.6 ± 0.7 s vs. 1.5% DSS, 8.8 ± 2.2 s, t = 3.22, df = 4, *p.adj* = 0.097; Male: 0% DSS, 6.6 ± 0.5 s vs. 1.5% DSS, 1.9 ± 0.5 s, t = 6.05, df = 4, *p.adj* = 0.011). Similarly, the number of burrows dug decreased over time (F (4, 96) = 17.301, *p* < 0.0001; Fig. 5E) and was found to be dependent on the dose of DSS that mice received (Interaction Dose:Day: F (8, 96) = 5.859, *p* < 0.001; Fig. 5E) as well as sex (Interaction Sex:Day: F(4, 96) = 3.053, *p* = 0.02; Fig. 5E). To confirm that the observed deficits in digging behavior were not the result of mice being physically incapacitated, mice were also assessed on the rotarod after each test of digging. No deficit in locomotor function was seen during the study (F (4, 143) = 1.543, *p* = 0.193; Fig. S4B).

A high co-incidence of inflammatory bowel disease and anxiety is reported in humans [3], and mice undergoing the DSS model of colitis exhibit increased anxiety behaviors [16]. In agreement, among respondents of the Impact of Chronic Pain on Daily Life survey diagnosed with a single condition, 71.43% of those living with inflammatory bowel disease strongly agreed that the pain caused by their diagnosis had an influence on their decision to leave the house, second only to those living with endometriosis (Fig. S4C). To further explore if this aspect of the human visceral pain could be resolved from the digging assay automated positional tracking was employed to quantify the explorative nature of mice in DSS colitis experiments (Fig. S4Di). Agreeing with human patients, mice experiencing colitis explored a smaller area of the digging cage as colitis-like symptoms developed (F (2, 53.87) = 26.94, *p* = 0.0079; Fig. 5F, S4D), with a dependency on sex (F (1, 48.05) = 48.05, *p* = 0.0033; Fig. 5F) This finding further highlights the parallels between the behaviors captured with the digging assay and the experiences of human patients, allowing additional insight into how painful conditions affect spontaneous mouse behavior.

Disease activity indexes are useful for monitoring animal welfare during experimental colitis, however histopathological assessment of the affected tissue offers a more conclusive assessment of disease extent. After behavioral assessments on day 7, mice were sacrificed, and colons extracted. In line with previous reports [18,35] a reduced colon length was observed for mice that drank DSS-containing water (F (2,18) = 15.947, *p* = 0.000104; Fig. S4E). Sections of the distal colon were prepared and stained (Fig. 5G), and histological scores were assigned. The dose of DSS that animals received had a significant influence over the resulting histological score (F (2, 18) = 48.089, *p* < 0.0001; Fig. 5H) and, in line with the DAI and time spent digging, a sex effect was also seen (F (1, 18) = 13.437, *p* = 0.002; Fig. 5H). Having confirmed the extent of intestinal inflammation on Day 7 for each mouse, histological scores were next compared to the digging activity (time spent digging) observed on the final day of the experiment, offering the most comprehensive assessment of whether digging activity is predictive of disease severity. A negative correlation was observed (β = -0.7793, p < 0.0001; R^2^ = 0.645, F (1, 22) = 39.97, *p* < 0.0001; Fig. 5I). Although, DAI is less conclusive, the relationship between DAI and amount of time mice spent digging was also explored, also showing a negative correlation (β = -0.7222, p < 0.0001; R^2^ = 0.377, F (1, 51) = 30.86, *p* < 0.0001; Fig. S4F). The findings described here thus strengthen the notion of digging as a valuable read out of disease severity.

**Figure 5.**
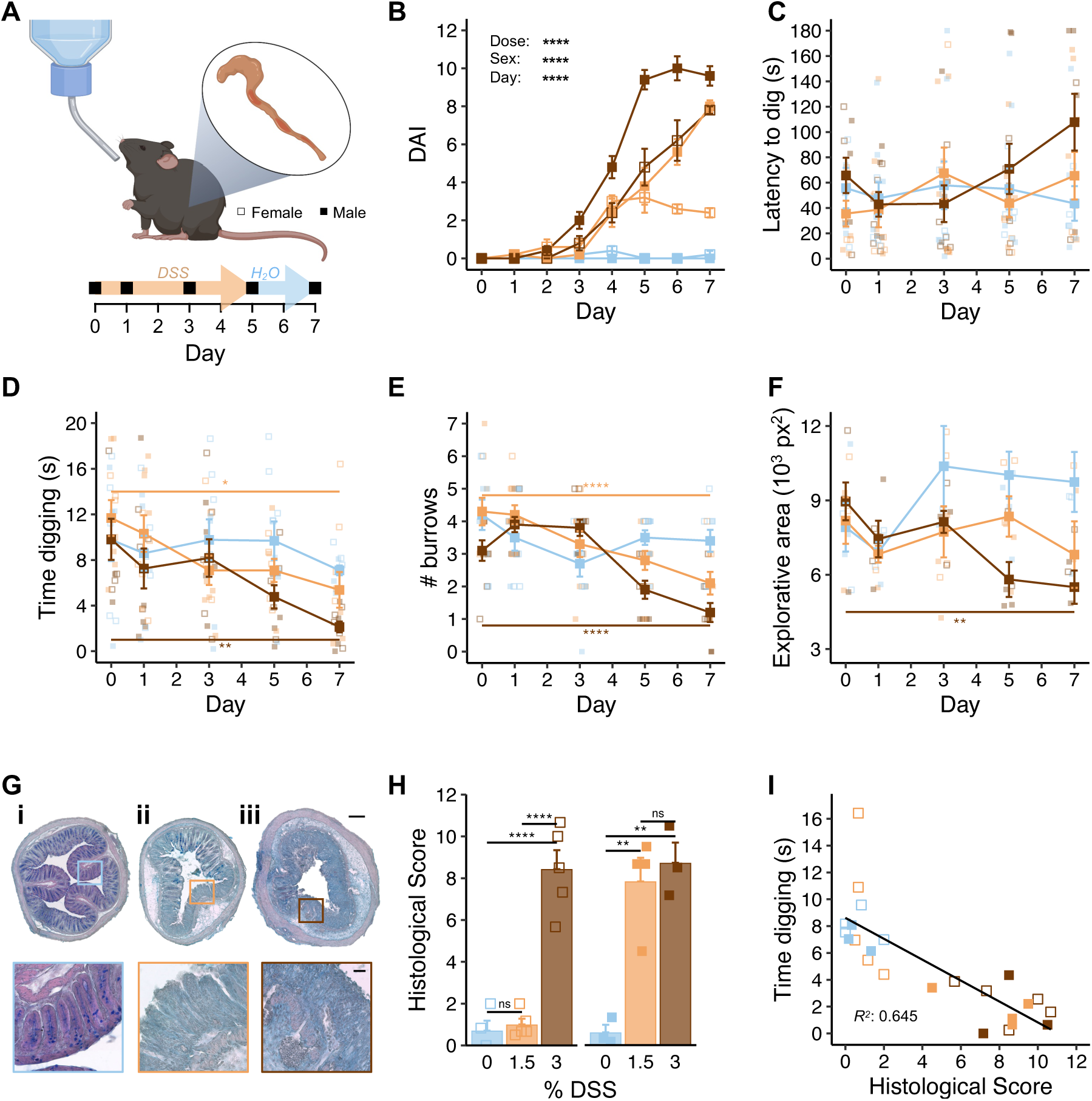
Digging behavior is predictive of the extent of intestinal inflammation in a visceral pain model. **(A)** Schematic representation of the DSS model (created with BioRender.com) and experimental timeline. **(B)** Disease activity index (DAI) over experimental time. The **(C)** latency to dig, **(D)** time spent digging and **(E)** number of burrows dug across experimental timeline. **(F)** Positional tracking was used to determine the area of the digging cage explored by mice. **(G)** Representative staining of colon sections from male mice that received **(i)** water only, **(ii)** 1.5% w/v DSS, **(iii)** 3% w/v DSS. Scale bars: Full cross-sections = 400 µm; inserts = 200 µm. **(H)** Histological scores of colon sections according to animal sex and treatment group. **(I)** Correlation between histological score and digging duration of mice as tested on Day 7. ∗ p < 0.05, ∗∗ p < 0.01, ∗∗∗∗ p < 0.0001: **(B – F)** Repeated measures ANOVA followed by Bonferroni corrected post-hoc, annotated statistical differences are the effect of time for each experimental group. **(H)** Two-way ANOVA followed by Bonferroni corrected post-hoc. **(I)** Simple linear regression. **(B-F)** All groups: n = 10, 5 females (denoted by open symbols) and 5 males (denoted by closed symbols). **(G-H)** 0% group: n = 4 females, 3 males; 1.5% group: n = 5 females, 4 males; 3% group: n = 5 females, 3 males.

## 4. Discussion

Animal models of painful conditions are an incredibly useful aspect of the drug discovery process, not just for pre-clinical testing of potential new pain medications, but also in the identification of putative targets. With the ever pressing need to deliver safer and more efficacious analgesics, ensuring that data obtained from animal models of painful conditions are as reliable and informative as possible is a top priority. Here we add to the mounting evidence that chronic pain patients are prevented from going about their everyday lives by the pain they experience (Fig. 1). We also show that an analogous everyday behavior of mice, digging, is negatively affected by several models of painful conditions (Figs 3 – 5). We thus believe there is strong justification to include digging as an ethological behavioral readout of animal wellbeing in the field of pain research. We believe that the addition of digging to behavioral characterization, will complement data obtained via other evoked and non-evoked assays, to generate a more comprehensive pain profile, which will aid in smoothing the transition from treating pain in animals to helping human patients.

Burrowing is the main alternative ethological rodent behavior to digging. Undeniably useful as a measure of animal wellbeing, deficits in burrowing behavior have been reported in inflammatory, visceral, neuropathic and post-surgical pain models [11,21,23,32,36,45]. However, we believe that digging [12] provides a better ethological readout than burrowing. Our approach to studying digging is reminiscent of the open field test, whereby mice are placed in a completely uniform environment, which they can freely explore: naturally inquisitive, this often involves digging. Experimental set ups for assaying burrowing involve providing animals with apparatus to burrow from, and can thus be considered task based, which raises questions over the motivation of animals to perform the behavior, especially as in many cases burrows are filled with food pellets. This is evident from observations that mice will spend more time exploring a designated burrow (plastic tube) than in the open when a burrow is available [39]. Motivation has a large sway over any animal’s behavior, but the digging assay removes the confounder of object exploration from the motivation to perform the measured behavior, but still allows motivation to be inferred. This is evident from our observation that mice retain full locomotive ability when placed on the rotarod, and as such are given no choice but to move, in all the pain models we explored. However, when tested in our digging paradigm, we report a reduction in behavior across all models tested. We therefore assume that mice also retain the ability to dig, but on some level make a conscious decision not to, and thus believe that digging represents a better way to capture both the motivation and ability of mice to engage in a routine behavior. Further evidence to support this comes from our observation of reduced explorative behavior in mice experiencing visceral inflammation, when positional tracking was used to quantify the area mice explore during digging testing.

Visceral pain conditions are among the worst managed by currently approved medications, evident from findings that anti-inflammatories often exacerbate intestinal inflammation [6,26] and opioid medication producing constipation as a common side effect. Assessment of visceral pain in lab animals is also limited, which may contribute to the lack of effective pharmaceutical treatments. Currently, visceromotor responses to colorectal distention, measured by electromyography, afford the most specificity in assessing visceral pain in rodents. However, experiments often involve invasive surgeries, evoked stimulation, anesthesia and restraint [25], a stark contrast to how visceral pain manifests, and is assessed, in humans. Here we have shown a negative effect on the digging activity of mice in an experimental model of colitis (Fig. 5). Our findings that digging deficits were graded in line with the dose of DSS animals received, and a correlation between digging activity on the final day of behavioral testing and histopathological analyses of disease extent suggest that digging behavior is predictive of internal inflammatory state and thus holds high value in assessing animal wellbeing in DSS colitis. This linkage of internal inflammatory state to digging behavior provides further support for digging analysis to be widely used in the behavioral arsenal of pain scientists, particularly those researching visceral conditions. It will be interesting to see how digging behaviors are affected by other models of visceral conditions associated with pain. Bashir and colleagues [4] have recently shown that mice undergoing a model of endometriosis are slower to burrow, from a tube filled with food pellets, than sham animals, however, both groups successfully clear the burrow within 24-hours. Such findings align with our argument that the burrowing assay is more task driven than digging. Studying how the digging behavior of mice is affected by this endometriosis model may allow for a more objective measure of animal welfare in the absence of an overt challenge.

We have previously demonstrated that analgesia, evident from an increase in digging activity, can be obtained in the acute CFA-induced knee inflammation model using a blocker of the transient receptor potential vanilloid 1 (TRPV1) ion channel and chemogenetic approaches [8,9]. We have also observed improved digging in the more chronic destabilization of the medial meniscus model of OA following intra-articular mesenchymal stem cell derived extracellular vesicle (MSC-EV) administration [1]. Here, we report a difference in the analgesic efficacy of gabapentin and meloxicam when focusing on their effects on digging behavior deficits in the MIA OA model. Despite both drugs remedying mechanical hypersensitivity in an evoked behavioral assay, gabapentin conferred minimal improvement in digging behavior, while meloxicam brought digging durations more in line with baseline behavior (Fig. 4G). Thus, digging behavior also holds value in the testing of analgesics, in addition to tracking welfare through pain models. Furthermore, we have shown that the ethological digging assay may lend itself better to assess analgesia than traditional evoked nociception assays. This is demonstrated by two findings of this study: 1) that meloxicam provides better analgesia than gabapentin, when measured in terms of its effect on digging behavior in an experimental model of osteoarthritis (Fig. 4G), and 2) that human OA patients who take similar medications to manage their pain are more satisfied with the pain relief afforded by NSAIDs than gabapentinoids (Fig. 4I-L). Of course, the utility of digging as a predictor of analgesic success could be further tested by expanding investigations to other analgesics already approved to treat conditions that can be modelled in mice and comparing how changes in digging align with patient opinion.

Here, we describe the use of digging as a behavioral read out of animal welfare in various pain models. The utility of digging has been demonstrated by the robustness of behavior in naïve mice (Fig. 2) and deficits observed following acute inflammation (Fig. 3) and more translationally relevant models of osteoarthritis (Fig. 4) and colitis (Fig. 5). Sufficient sensitivity of the assay also allowed capture of graded responses, which correlated with the extent of inflammation. It is important to note the parallels between the observed deficits in digging behavior seen in mouse models of painful conditions and the reduced motivation and/or ability of humans experiencing pain to undertake routine behaviors, as well as the limitations of gabapentin on improving arthritis-triggered digging deficits in mice and complaints regarding the effectiveness of analgesia achieved when similar drugs are used by humans. Together, these demonstrate that digging is a suitable behavior for assessing pain and analgesia in mice. Indeed, the similarities in digging deficits and the human experience of pain potentially permit greater insight into the analgesic potential of novel drugs by measuring their ability to improve the likelihood of engaging in natural behavior. Furthermore, digging behavior can be assessed with minimal investment in equipment or personnel training, animals can be tested in batches and analyses performed offline after blinding. The nature of data collection is also highly amenable to automated analysis, indeed in future work we seek to expand the positional tracking performed here to train models capable of identifying spontaneous behaviors, including digging. Although we have previously shown that regulating articular neuron activity improves digging behavior in acute [8] and chronic [1] joint pain models, we cannot say with certainty that observed deficits in digging behavior are a result of animals experiencing pain. However, we have also argued that our approach allows a more objective assessment of the motivation to spontaneously perform an ethological behavior than alternatives. In this way, we believe that digging adeptly captures a critical aspect of the current IASP definition of pain [41], owing to its high reliance on emotional experience. This is different to many other behavioral measures of pain, which have a higher degree of affective reaction to passive overt or evoked stimuli, as opposed to digging being driven, mostly, by free will. That said, we are advocates for using multiple approaches to profile pain, agreeing with the recent recommendation of Sadler, Mogil and Stucky [42] to increase diversity and resolution of pain behavior measurements, believing that digging has certainly proved its use to become more commonplace.

## Acknowledgements

The authors thank the following charities for their assistance in promoting and recruiting pain patients to ‘The Impact of Chronic Pain on Daily Life’ survey: Arthritis Action, Arthritis Australia, Arthritis New Zealand, Arthritis Society Canada, Bowel Research UK, Crohn’s and Colitis Canada, Crohn’s and Colitis New Zealand, Crohn’s and Colitis UK, Endometriosis New Zealand, Endometriosis UK, Fibromyalgia Action UK and National Arthritis Foundation Singapore. The authors also thank the staff of the University of Cambridge Combined Animal Facility for their work in supporting this research. For the purpose of open access, the authors have applied a Creative Commons Attribution (CC BY) licence to any Author Accepted Manuscript version arising from this submission.

LAP and ESS acknowledge funding from the MRC (MR/W002426/1). AMC, GC and ESS were funded by Versus Arthritis (RG21973). GC and ESS were funded by the Rosetrees Trust (CM818). SC was supported by a Gates Cambridge Trust scholarship. HH was supported by a BBSRC/GSK iCASE PhD studentship (BB/V509528/1). RHR, MM, LWP and AR were supported by AstraZeneca PhD Studentships (RHR: G104108; MM: G104109; LWP: G113502; AR: G115018). MD and ESS acknowledge funding from the Wellcome Trust (225856/Z/22/Z). JPH and DCB acknowledge funding from Crohn’s and Colitis UK (PC2019/1-Bulmer).

## Author contributions

conceptualization: L. A. Pattison, S. Chakrabarti and E. St. J. Smith; investigation: L. A. Pattison, A. Cloake, S. Chakrabarti, H. Hilton, R. H. Rickman, J. P. Higham, M. Y. Meng, L. W. Paine and G. Callejo; data analysis: L. A. Pattison, A. Cloake, S. Chakrabarti, H. Hilton, R. H. Rickman, J. P. Higham, M. Y. Meng, L. W. Paine, M. Dannawi, L. Qui and A. Rittoux; writing: L. A. Pattison and E. St. J. Smith; supervision: D. C. Bulmer and E. St. J. Smith; funding acquisition: G. Callejo, D. C. Bulmer and E. St. J. Smith.

## Availability of data

Behavioral data sets supporting the conclusions of this article are available in University of Cambridge Apollo Repository (https://doi.org/10.17863/CAM.100117). Summary data from The Impact of Chronic Pain on Daily Life survey will be shared on reasonable request from the corresponding authors.

