## Supplementary Figures and Tables for "Digging deeper into pain – an ethological behavior assay correlating well-being in mice with human pain experience"

Figure S1

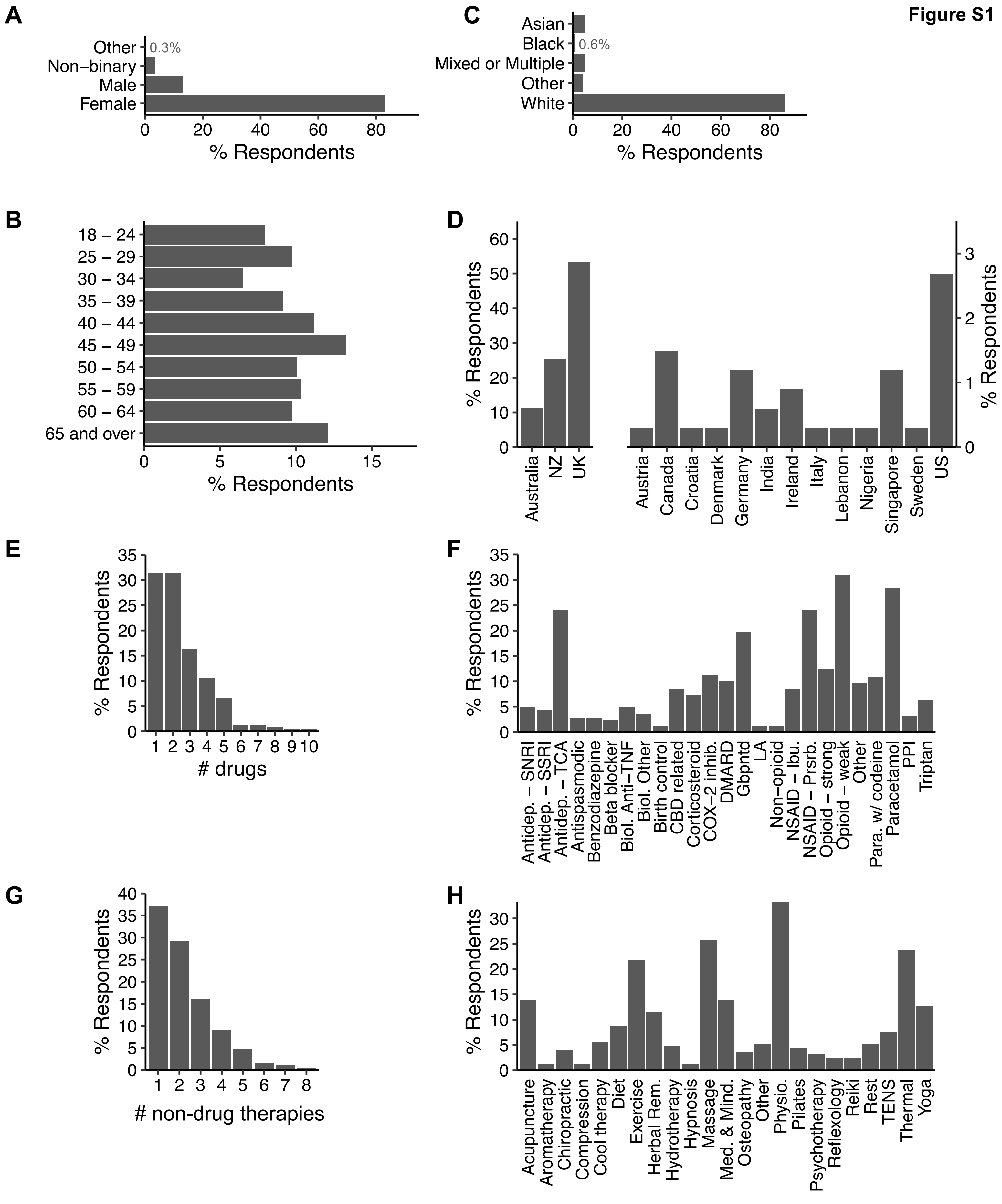


**Figure S1. Survey participant demographics, use of drugs and non-pharmacological therapies to manage pain.** Demographical breakdown of participant **(A)** gender, **(B)** age, **(C)** ethnicity, **(D)** country of residence (NZ = New Zealand). **(E)** Frequency distribution of the number of drugs used by respondents. **(F)** Summary of drug types used by respondents to mange pain (Antidep. = antidepressant; BB = beta blocker; Biol. = biological therapy; CBD = cannabidiol; DMARD = disease modifying antirheumatic drug; LA = local anaesthetic; NSAID = non-steroidal anti-inflammatory drug; Ibu. = ibuprofen; Prsrb. = prescribed; Para. = paracetamol; PPI = proton-pump inhibitor). See Table S2 for a breakdown of 'Other' and "Other biological therapy' categories. **(G)** Frequency distribution of the number of non-drug therapies used by respondents. **(H)** Summary of non-drug therapies used by respondents to manage pain (Med. & Mind. = meditation & mindfulness; Physio = physiotherapy; TENS = transcutaneous electrical nerve stimulation). See Table S3 for a breakdown of 'Other' category.

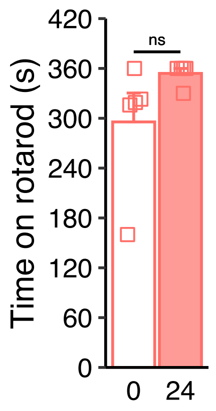


**Figure S2. Locomotor coordination is not affected by CFA-induced inflammation of the knee joint.** Mice were tested on a rotarod before and 24-hours post-injection of CFA to one knee joint. Wilcoxon signed-rank test. n = 5 females.

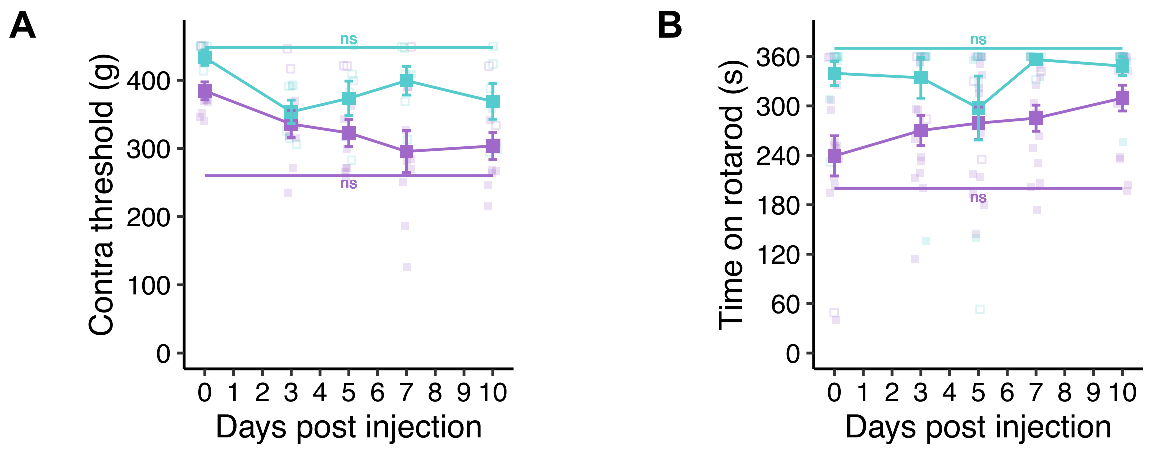


**Figure S3. Mechanical threshold of the contralateral knee and locomotor coordination are not affected by MIA model of osteoarthritis.** Mice were tested on a rotarod before and following injection of MIA to one knee joint. Repeated measures ANOVA. **(A)** n = 2 MIA females, 8 MIA males, 5 saline females, 1 saline male. **(B)** n = 2 MIA females, 13 MIA males, 5 saline females, 4 saline males. Females are denoted by open symbols and closed symbols represent males for individual data points.

**Figure S4**

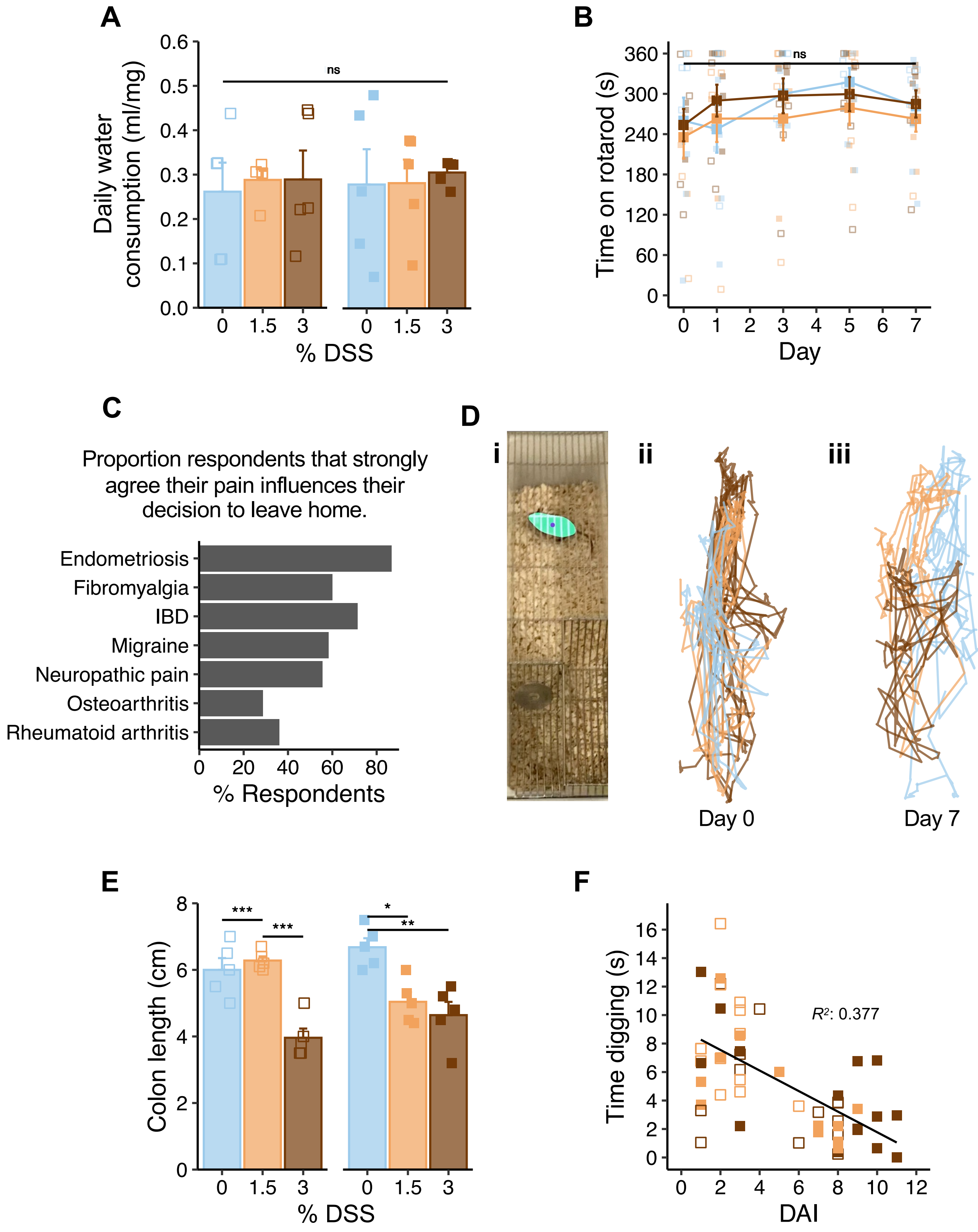


**Figure S4. Average daily water consumption, time on rotarod, locomotion, colon length and correlation between disease activity and digging time for mice on DSS model. (A)** Average daily water consumption, corrected for body weight, of mice from each treatment group by sex. **(B)** Average time on rotarod across experimental timeline. **(C)** Proportion of human patients with a single diagnosis of a condition that causes them chronic pain who strongly agree their pain affects their motivation to leave home according to diagnosis. **(D) (i)** Automated analyses were used to track the position of the center-point of mice from digging videos. Representative motion traces of male mice from each experimental group on **(ii)** day 0 and **(iii)** day 7. **(E)** Colon lengths at end of study, by treatment group and sex. **(F)** Correlation between disease activity index (DAI) and digging duration of mice on 3% w/v or 1.5% w/v DSS. $* \ p < 0.05$, $** \ p < 0.01$, $*** \ p < 0.001$: **(A,C)** Two-way ANOVA **(C)** followed by Bonferroni corrected post-hoc. **(B)** Three-way ANOVA. **(E)** Simple linear regression. n = 10 per group, 5 females (denoted by open symbols), 5 males (denoted by closed symbols).

| **Condition** | **# respondents** |
|---|---|
| Endometriosis | 40 |
| Fibromyalgia | 127 |
| IBD | 41 |
| IBS | 75 |
| Migraine | 76 |
| Neuropathic pain | 73 |
| Osteoarthritis | 88 |
| Other arthritic<br>*- Ankylosing spondylitis*<br>*- Early inflammatory arthritis*<br>*- Facial arthropathy*<br>*- Gout*<br>*- Inflammatory polyarthropathy*<br>*- Peripheral arthritis*<br>*- Psoriatic arthritis* | 47<br>*- 15*<br>*- 1*<br>*- 1*<br>*- 1*<br>*- 2*<br>*- 2*<br>*- 25* |
| Other musculoskeletal<br>*- Bursitis*<br>*- Carpal tunnel syndrome*<br>*- Connective tissue disorder*<br>*- Hypermobility syndrome*<br>*- Idiopathic musculoskeletal chronic pain*<br>*- Knee pain*<br>*- Osteoporosis*<br>*- Spinal / back condition*<br>*- Temporomandibular joint disfunction* | 33<br>*- 2*<br>*-1*<br>*- 2*<br>*- 9*<br>*- 1*<br>*- 2*<br>*- 1*<br>*- 13*<br>*- 2* |
| Other visceral<br>*- Adenomyosis*<br>*- Chronic pancreatitis*<br>*- Cystitis / painful bladder syndrome*<br>*- Focal segmental glomerulosclerosis*<br>*- Microscopic colitis*<br>*- Polycystic ovary syndrome*<br>*- Small intestinal bacterial overgrowth*<br>*- Vulvodynia* | 16<br>*- 1*<br>*- 1*<br>*- 8*<br>*- 1*<br>*- 1*<br>*- 2*<br>*- 1*<br>*- 1* |
| Other<br>*- Behçet's disease*<br>*- Chronic fatigue syndrome*<br>*- Conn's syndrome*<br>*- Da Costa syndrome*<br>*- Long covid*<br>*- Post sepsis syndrome*<br>*- Psychosomatic*<br>*- Raynaud's syndrome*<br>*- Sjögren's syndrome*<br>*- Systemic Lupus Erythematosus* | 14<br>*- 1*<br>*- 2*<br>*- 1*<br>*- 1*<br>*- 1*<br>*- 1*<br>*- 1*<br>*- 1*<br>*- 1*<br>*- 4* |
| Post-operative pain | 14 |
| Rheumatoid arthritis | 48 |

**Table S1. Full breakdown of diagnoses which cause survey participants chronic pain relating to Figure 1A.**

| Biol. Other | Other |
|---|---|
| Anti-CD20 (1)<br>Anti-CGRP (5)<br>Anti IL6R (1)<br>Anti-IL17 (2) | Angiotensin receptor blocker (2)<br>Anticonvulsant (3)<br>Antidiarrheal (1)<br>Antiemetic (1)<br>Antifibrinolytic (2)<br>Antihistamine (1)<br>Antihypertensive (1)<br>Antipsychotic (1)<br>Calcium channel blocker (1)<br>Integrin blocker (1)<br>Laxative (1)<br>Neurotoxin (1)<br>Opioid receptor antagonist (1)<br>Other anti-inflammatory (2)<br>Other antirheumatic (4)<br>Statin (1) |

**Table S2. Breakdown down of drugs classified as 'Other biologic' or 'Other' relating to Figure S1F.**

| Other |
|---|
| Reading (1)<br>Piercing (1)<br>Homeopathy (3)<br>Cognitive behavior therapy (1)<br>Podiatry (1)<br>Cupping (1)<br>Singing (1)<br>Distraction (3)<br>Outdoors (1) |

**Table S3. Breakdown down of non-drug therapies classified as 'Other' relating to Figure S1H.**
